# Circadian control of kidney regeneration via Lima1-mediated regulation of EMT

**DOI:** 10.1101/2024.04.02.587728

**Authors:** Xian He, Ziming Wang, Linxi Cheng, Han Wang, Yuhua Sun

**Affiliations:** Key Laboratory of Breeding Biotechnology and Sustainable Aquaculture, Institute of Hydrobiology, Chinese Academy of Sciences, Wuhan 430072, China; The Innovation of Seed Design, Chinese Academy of Sciences, Wuhan 430072, China; Hubei Hongshan Laboratory, Wuhan 430070, China; Soochow University, China

**Keywords:** circadian clock, cytoskeleton, renal regeneration, zebrafish

## Abstract

The circadian clock genes are known important for kidney development, maturation and physiological functions. However, whether and how they play a role in renal regeneration remain elusive. Here, by using the single cell RNA-sequencing (scRNA-seq) technology, we investigated the dynamic gene expression profiles and cell states after acute kidney injury (AKI) by gentamicin treatment in zebrafish. The core clock genes such as *per1/2* and *nr1d1*, which encode transcriptional repressors of the circadian system, are strongly induced in the proximal tubule epithelial cells (PTECs). By generating mutant zebrafish lines, we show that *per1* and *nr1d1* are required for proper renal regeneration, by facilitating the expression of renal progenitor cell (RPC) genes. In *per1* and *nr1d1* mutants, the expression of RPC genes and the number of RPCs were decreased, resulting in a marked delay in nephron regeneration. *lima1a*, which encodes a cytoskeleton binding protein that functions to negatively regulate epithelial to mesenchymal transition (EMT), is identified as the direct target of the clock proteins. Down-regulation of *lima1a* is associated with enhanced EMT, increased expression of RPC markers, and accelerated nephron regeneration. We propose that up-regulation of *per1* and *nr1d1* after AKI facilitates the formation of nephrongenic RPCs by repressing *lima1a.* Our findings using zebrafish provide important insights into the roles of the circadian clock genes in kidney repair.

Graphic abstract

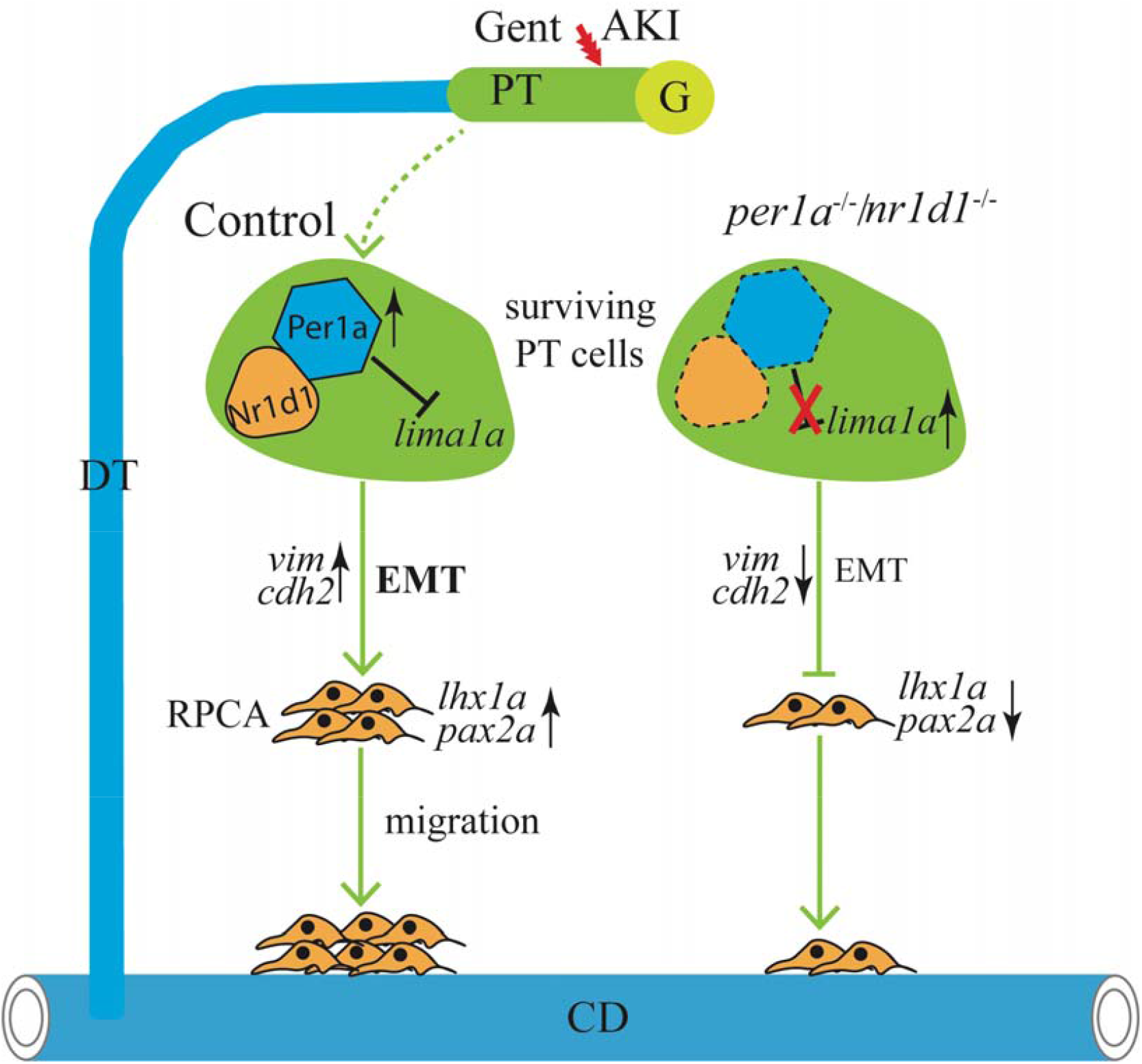

## Introduction

Kidney health and function can be affected by many factors, including excessive exposure to environmental pollutants or antibiotic medications. Acute kidney injury (AKI), a common and severe disease, may affect all segments of the nephron tubules, but the proximal tubule epithelial cells (PTECs) suffer the most injury due to its high metabolic activities.^1, 2^ Repairing the cells requires the dedifferentiation, proliferation, and migration of surviving PTECs to replace dead cells after AKI.^3–6^

Compared to mammals, zebrafish have a remarkable capacity to regenerate the kidneys. Zebrafish renal regeneration can be largely divided into 4 stages: First, the PTECs die by chemical or physical injuries, accompanied by early response and inflammation reaction; Second, the survived tubular epithelial cells (TECs) undergo a series of cellular events, including epithelial to mesenchymal transition (EMT),^7^ dedifferentiation, and metabolic reprogramming.^8, 9^ Third, renal progenitor cells (RPCs), marked by *lhx1a* and *wt1b*, proliferate and re-differentiate to repair the damaged nephrons *via* a mesenchymal to epithelial (MET) process.^3, 10–12^ Fourth, the renal regeneration is terminated after the restore of epithelial cell morphology and the nephron function. Despite the recent progress in the field, many outstanding questions remain to be answered. For instance, what are the upstream regulators or signal molecules for stimulating the proliferation and migration of *lhx1a^+^* RPCs? Fibroblast growth factors (Fgfs) and WNT, which are EMT inducers, have been shown to play major roles in formation and migration of RPCs.^3, 13^ Of note, Fgfs are induced specifically in TECs at 3 dpi, when dedifferentiation- and mesenchymal markers such as *vimentin* (*vim*) are predominantly expressed.^3^ These observations raise a possibility that injured TECs acquire stem cell-like properties during dedifferentiation and EMT, then migrate to regenerate the nephrons.^14, 15^

*LIM domain and actin-binding protein 1* (*LIMA1*), also known as *Epithelial protein lost in neoplasm* (EPLIN), encodes an actin-binding cytoskeletal protein containing a LIM structural domain.^16, 17^ Lima1*/*Eplin is a well-known cytoskeletal regulator, which links the cadherin-catenin complex to the actin cytoskeleton, and enhances the bundling of actin filaments by inhibiting actin filament depolymerization. In mice, *Lima1* is shown to be expressed in renal PTECs and glomeruli cells, and modulate cell adhesion and movement, likely regulated by FGF/ERK mediated phosphorylation.^18, 19^ In human, Lima1 has been identified as one of the hub genes that are linked with kidney disease.^20^ In fact, *Lima1* is a negative regulator of EMT,^21–23^ an essential process in kidney regeneration.^24^

The kidney is one of the top organs that display rhythmically expressed genes in vertebrates. Many of renal functions, including electrolyte excretion, blood flow, and filtration, exhibit circadian rhythms, and under circadian regulation,^25–30^ and dysfunction of circadian clock can lead to various renal disease conditions.^31^ Although the clock genes such as *Per2* have been shown to regulate mouse embryonic kidney development,^32^ its role in kidney regeneration has not been well defined.^33^ In this work, we surprisingly found that genes encoding transcriptional repressors of the circadian clock, including *per1/2* and *nr1d1*, were induced by AKI, in particular in TECs. By generating circadian clock gene mutants of zebrafish, we provided compelling evidence that *per1* and *nr1d1* are required for proper nephron regeneration through regulation of actin cytoskeleton via repressing *lima1a*.

## METHODS

Detailed description of experimental design and methods can be found in the Supplementary Methods.

### Zebrafish maintenance

Zebrafish were obtained from the China Zebrafish Resource Center, Institute of Hydrobiology, Wuhan, China. Fish were maintained in a 12h light/12h dark cycle.

### Generation of mutant zebrafish lines by CRISPR/Cas9

The zebrafish mutant alleles were generated by CRISPR/Cas9 system. Two guide RNA were designed on exon2 and exon5 for the *nr1d1* gene, and the gRNA sequences were CCCAACCGTACCAGCCCTGTG, CCTCACCGGCTCCAACCTCCCC, respectively. gRNAs sequences for *per1a* and *per1b* were CCTCAACCTGTAGCTCACTGC and CCATGGGTATGGAGACAACGG, respectively. One guide RNA was designed for the exon 2 of the *lima1a* gene, and the sequences were CCCTCATCGAAAAGCCCACCG. 200 pg purified guide RNAs were mixed with 250 pg Cas9 protein (Invitrogen, A36498), and were injected into 1-cell stage embryos.

### Acute kidney injury by genetamicin treatment

To induce acute kidney injury, 5 month-old adult wild type and mutant zebrafish (average weight of 0.5g) were injected with gentamicin (80 mg/kg body weight) at 10: 00-11: 00 am. Similar volume of PBS was injected, and served as control. The kidneys were peeled from anesthetic animals at 1, 3, 5, 7, 10 dpi. At least 3 experimental repeats were performed. The whole kidneys from 8-10 animals per group or its sections were used for qRT-PCR, IF, and HE staining analyses.

### Bulk RNA-sequencing

8-10 kidneys were pooled for each time point for bulk RNA-sequencing. Total RNA were isolated using the TRIzol reagent (ThermoScientific, USA). RNA-seq experiment has been performed in at least two replicates. The pooled RNA samples were sent to the BGI company, China, where RNA-sequencing libraries were constructed and sequenced by a BGI-500 system. All the RT-PCR primers can be found in the Supplementary file 1.

Differentially expressed genes (DEGs) were defined by FDRLJ<LJ0.05 and a Log2 fold change> 1. For heatmap analysis, TPM were used and the plot was made by pheatmap. Gene ontology (GO) analysis for differentially expressed genes (DEGs) were performed at https://geneontology.org (accessed on 23-November-2022).

### Single cell RNA-sequencing analysis

Single cell RNA-sequencing was performed by the Gene Denovo company, Guangzhou, China. The detailed information of data analysis can be found in the Supplementary methods.

### Whole mount in situ hybridization (WISH)

The RNA probes were used to detect the expression of *slc20a1a, trpm7, slc12a1, slc12a3, lhx1a, wt1b pax2a, irx3b, sox9a, sall1a, nr1d1, nr1d2a, per1a, per1b, atf3, egr1, egr4,* and *lima1a*. Primers used for probe synthesis were provided in the supplementary file 1.

### Statistical analysis

Statistical analysis was carried out using Student’s *t* test to compare differences. All statistics were done using Prism 5.0 software (Graphpad Software Inc). An analysis of variance (ANOVA) was used to test the statistical significance, when multiple comparisons were conducted. Results are shown in the figure legends or in the figures.

## RESULTS

### scRNA analysis of kidney cells

To induce the AKI, 5-month old zebrafish were intraperitoneally injected with 80 ng/mg of gentamicin, an established nephrotoxin.^34^ Nephron damage was evidenced at 1 dpi (day post injection), by white casts of dead epithelial tissue excreted by the injured fish, loss of expression of the proximal tubule (PT) marker *slc20a1a*, and failure to uptake the fluorescent dextran (Supplementary Figures 1A-D).

To understand the mechanisms underlying renal regeneration, non-injured and injured kidneys at 1, 3, and 5 dpi were subjected to single cell RNA-sequencing (scRNA-seq) (Supplementary file 2). These time points were selected in order to explore the early molecular and cellular events that are critical for kidney repair (Figure 1A). First, we analyzed the scRNA-seq data from the non-injured kidneys. After quality control filtering, a total of 18,361 cells (n=2) were obtained, and 1500 genes were detected in each single cell on average. Unsupervised clustering analysis was performed, and the Uniform Manifold Approximation and Projection (UMAP) revealed a total of 16 major clusters (Figure 1B). Based on the cell type specific markers, the identity of the individual clusters was determined (Figure 1C). Kidney cells can be grouped to 7 major cell lineages: *nphs2* expressing podocytes, *slc12a1/slc20a1a* expressing nephron epithelium, *gata3* expressing pronephric duct cells, *mpx* and *c1q* expressing immune-related cells, *itln1* expressing mucin cells, *cdh5* and *sele* expressing vascular endothelium, and *col1a2* expressing stromal/fibroblast cells.^35^ The nephron epithelium can be further divided into 3 clusters: (a) the proximal tubule cells (PTECs), defined by the expression of *slc20a1a, slc13a1* and *lrp2a*, (b) the distal tubule cells (DTECs), marked by *slc12a1* and *slc12a3,* and (c) the nephron epithelium expressing *cldnh* and *tnfrsf11b*.

**Figure 1.**
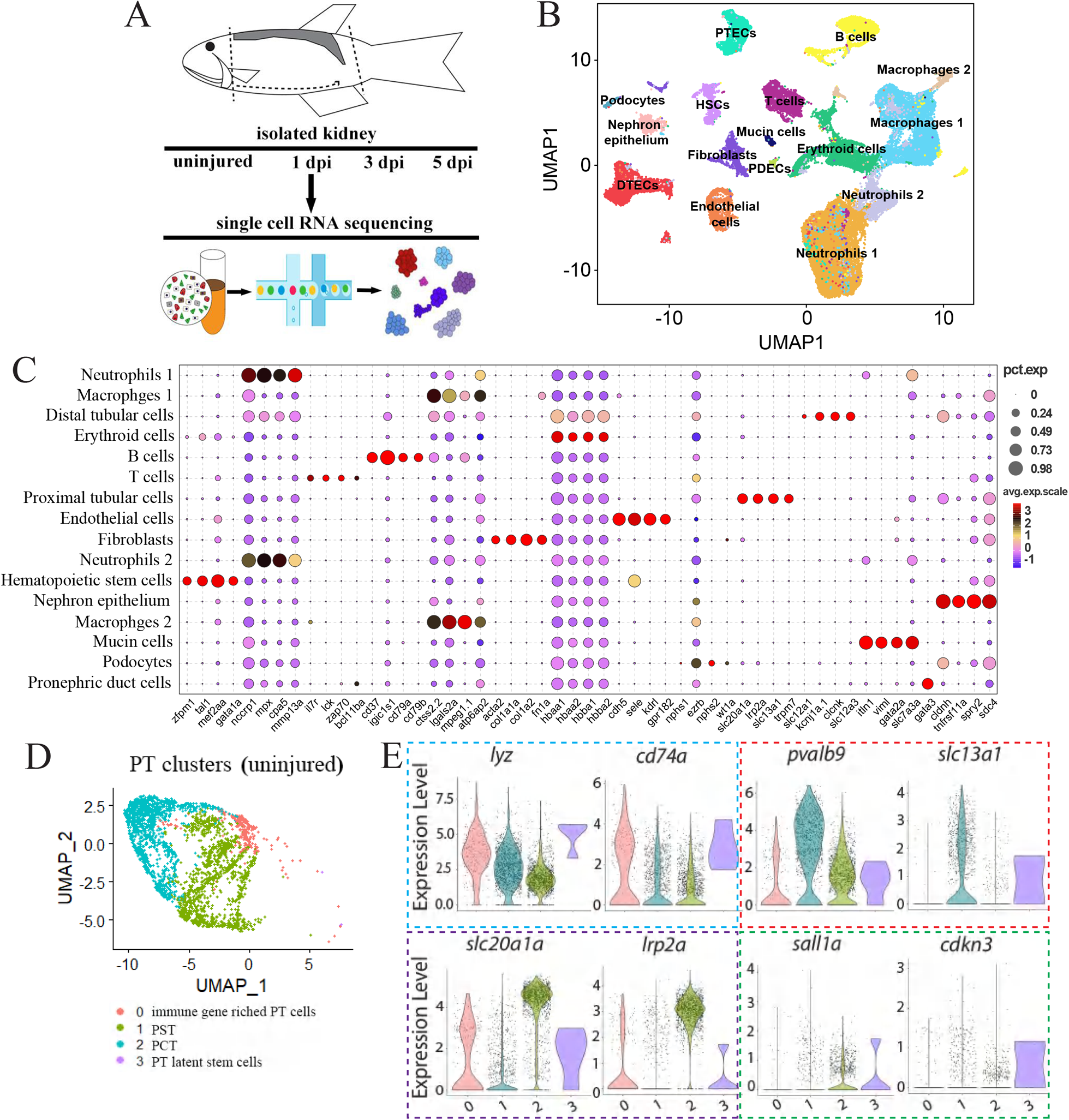
scRNA-seq data analysis of zebrafish kidneys. (**a)** Schematics of the study design. (**b)** UMAP embedding of scRNA-seq data. Using kidney marker genes, cells were annotated into the indicated 16 major clusters. PTECs: proximal tubular epithelial cells; DTECs: distal tubular epithelial cells; B cells: B lymphocytes; T cells: T lymphocytes; PDECs: pronephric duct cells; PD; podocytes; HSCs: hematopoietic stem cells. (**c)** Dot plots displaying cluster-enriched marker genes. (**d)** UMAP projection of PT cells, demonstrating 4 sub-clusters. The subcluster 0: the immune gene enriched population; the subcluster 1: PST; the subcluster 2: PCT; the sub-cluster 3: the resident stem cell population. (**e)** Violin plots grouped by meta-clusters, demonstrating the subcluster-specific expression of the indicated genes. The dashed colored boxes were used to indicate the markers of the 4 subclusters. 0-3 of the x-axis: the subclusters 0-3.

We then analyzed the scRNA-seq data from the injured kidneys at different time points. After quality control filtering, a total of 16,755 cells from 1 dpi kidneys; 21,565 cells from 3 dpi kidneys; and 17,745 cells from 5 dpi kidneys were obtained. An average of 1500 genes were detected in each single cell, indicating of high quality of our scRNA data. Analysis of UMAP showed that there was no cluster gain or loss by gentamicin treatment. Nevertheless, a dramatic change in number of PTECs was observed (Supplementary table 1), in line with that they suffer the most of injury.

A pseudo-time analysis showed that the PT cluster was at the start of the trajectory, supporting that PTECs contribute to renal regeneration (Supplementary Figures 1E). We focused on the PT cluster (n=1327) and performed the second-level clustering in uninjured kidneys, which resulted in 4 subclusters (Figure 1D; Supplementary Figure 1F; Supplementary file 3). The subcluster 0 (n=164) was enriched for immune genes such as *cd74a* and *lyz*, indicative of a PT population with immune cell characteristics (Figure 1E). The subcluster 1 (n=529) was identified as the proximal straight tubule (PST) as it abundantly expressed PST markers *slc13a1* and *pvalb9*. The subcluster 2 (n=630) was identified as the proximal convoluted tubule (PCT) marked by *slc20a1a*, *slc34a1a* and *lrp2a.* The subcluster 3 (n=4) was consisted of a small number of cells that expressed renal stem/progenitor cell markers such as *sall1* and *pax2a*.^1, 36–39^ This subcluster also expressed cell cycle inhibitor genes such as *cdkn1/3,* suggesting that the cell type was in a latent state in uninjured condition. Based on the gene expression signature, we propose that the subcluster 3 might be a latent stem cell population.^40, 41^

### Dynamic gene expression of 4 PT subclusters

To understand how the surviving PTECs contribute to renal regeneration, we characterized the cellular and molecular dynamics of individual subclusters (Figures 2A-B). At 1 dpi, there was a huge loss of PST and PCT cell types, accompanied by a significant peak of cell proliferation of the immune gene enriched sub-population (the subcluster 0) (Figures 2B; Supplementary Table 2). Induction of early response- and stress- (*atf3*, *egr1/3*, *nfkb*, and *jun*) genes was observed, highlighting the immediate response of kidney to gentamicin (Figure 2C-D; Supplementary Figure 2A-C). Some glycolytic-related genes such as *pdk2/4* and *pkma* were up-regulated, which was in line with that there is a metabolic switch (from oxidative phosphorylation to glycolysis) during early stages of renal regeneration.^42^ Moreover, there was an increased expression of transcription factor genes that are normally expressed during embryonic kidney development, including *sox9a* and *emx1*,^43, 44^ indicating that the surviving PTECs were undergoing dedifferentiation and resembled an “embryonic form” on a transcriptomic level. Interestingly, core clock genes such as *per1/2* and *nr1d1*, which encode transcriptional repressors in the circadian system, were induced. By contrast, *bmal1* and *clock1*, which encode transcriptional activators, remained lowly expressed (Figure 2C-D).

**Figure 2.**
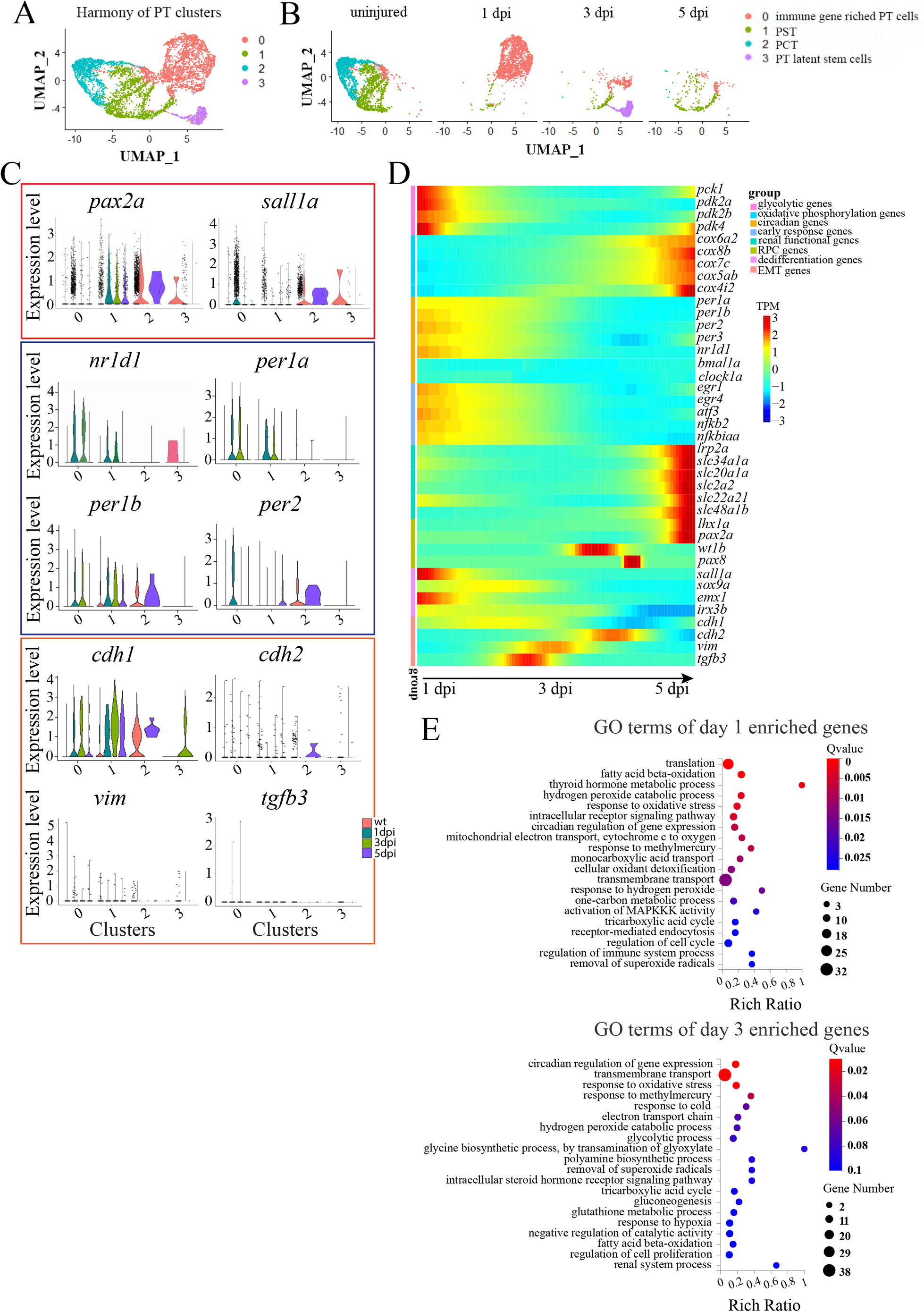
Dynamic gene expression of the 4 PT subclusters during renal regeneration. (**a)** UMAP embedding of all cells profiled during early stages of renal regeneration. (**b)** UMAP embedding of scRNA-seq data colored by time points highlights the progressive nature of 4 subclusters during regeneration. (**c)** Violin plots showing dynamic expression of the indicated genes, grouped into markers of RPCs, circadian clock and EMT. The colored boxes were used to highlight the indicated progenitor-, clock-, and EMT-genes. 0-3 of the x-axis: the subclusters 0-3. (**d)** Heat map showing the dynamic expression of the indicated genes, grouped into 8 categories that are shown on the upper right corner. (**e)** GO terms of the induced genes at 1 and 3 dpi.

At 3 dpi, a robust cell proliferation of the subcluster 3 was observed, accompanied by increased expression of *pcna*, a cell proliferation marker and *havcr1/kim1*, a renal injury and repair marker (Supplementary Figure 2C), suggesting that the latent stem cells went from quiescent to activation state and participated in renal repair. Meanwhile, the expression of mesenchymal markers such as *cdh2/N-cad* and *vim* was increased (concomitant with decreased expression of *cdh1/E-cad*), indicative of EMT in the surviving PTECs (Figure 2C-D). At 5 dpi, co-expression of RPC markers (*lhx1a* and *wt1b*) and renal transporter genes (the *slc* family members) was observed, suggesting that intrinsic TECs contribute to nephron repair by becoming *lhx1a^+^* RPCs. Consistently, GO (gene ontology) enrichment analysis of day 1 up-regulated genes showed terms such as circadian regulation of gene expression, response to stress, and regulation of immune system, and GO analysis of day 3/5 up-regulated genes showed terms such as regulation of cell proliferation, regulation of epithelial development, transmembrane and sodium ion transport, and renal system process (Figures 2E; Supplementary Figure 2D).

Whole mount in situ hybridization (WISH) was performed to verify the expression of gentamicin-induced genes (Figure 3A-D; Supplementary Figure 3A). In uninjured control kidneys, these genes were barely expressed. After gentamicin treatment, they were induced, exhibiting peaks at different time points. To substantiate this, we performed bulk RNA-sequencing for kidneys at 1, 3, 5, 7 dpi (Supplementary file 4). The results were consistent with that revealed by our scRNA analysis (Figure 3E; Supplementary Figure 3B).

**Figure 3.**
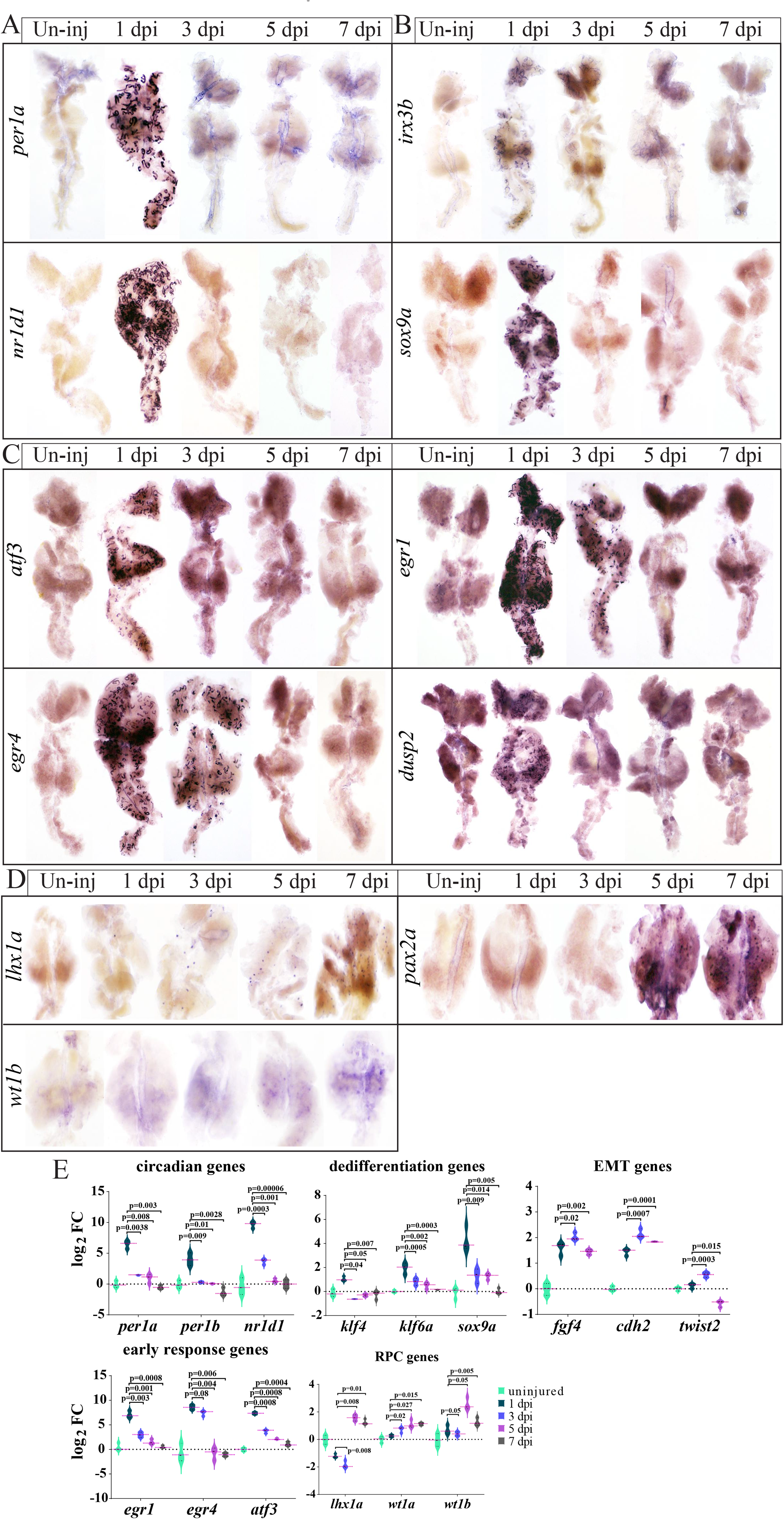
Validation of the dynamic expression of the indicated genes. (**a)** WISH images of the clock genes. (**b)** WISH images of the dedifferentiation genes. (**c)** WISH images of the early response genes. (**d)** WISH images of the RPC genes. (**e)** Plots showing the dynamic expression of the indicated genes during a time course of 7 days after AKI, based on 3 bulk RNA-seq replicates. Log2 FC: the log2 fold change of the indicated gene’s expression at the indicated time points relative to day 0 (uninjured kidneys). P-values were indicated to show statistical significance. WISH were repeated three times, and shown are representative data.

The above results confirmed the coordinated and progressive nature of renal regeneration,^2^ but additionally highlighted two key observations: the circadian clock genes are induced in PTECs at 1 dpi, and RPC markers are expressed within PTECs at 5 dpi. The latter observation is in accordance with the notion that intrinsic TECs are responsible for nephron repair,^42, 45^ and the former implied that the clock genes are important for nephron regeneration.

### *per1a* and *nr1d1* are required for proper renal regeneration

We examined every 4h for the expression of the core clock genes in zebrafish kidneys before/after gentamicin treatment. In kidneys of zebrafish injected with PBS, the core clock genes were rhythmically expressed (Figure 4A). *bmal1a* and *clock1a* displayed peaks at ZT12 and troughs at ZT0, respectively. *per1* and *nr1d1* were moderately expressed, showing antiphase patterns: *per1a* displayed a peak and a trough at ZT8 and ZT20, respectively, whereas *nr1d1* displayed a peak and a trough at ZT20 and ZT8, respectively. After gentamicin treatment, *per1a, per2* and *nr1d1* were significantly induced (*P*< 0.05). By contrast, the expression of *bmal1a* and *clock1a* was barely altered (Figure 4B).

**Figure 4.**
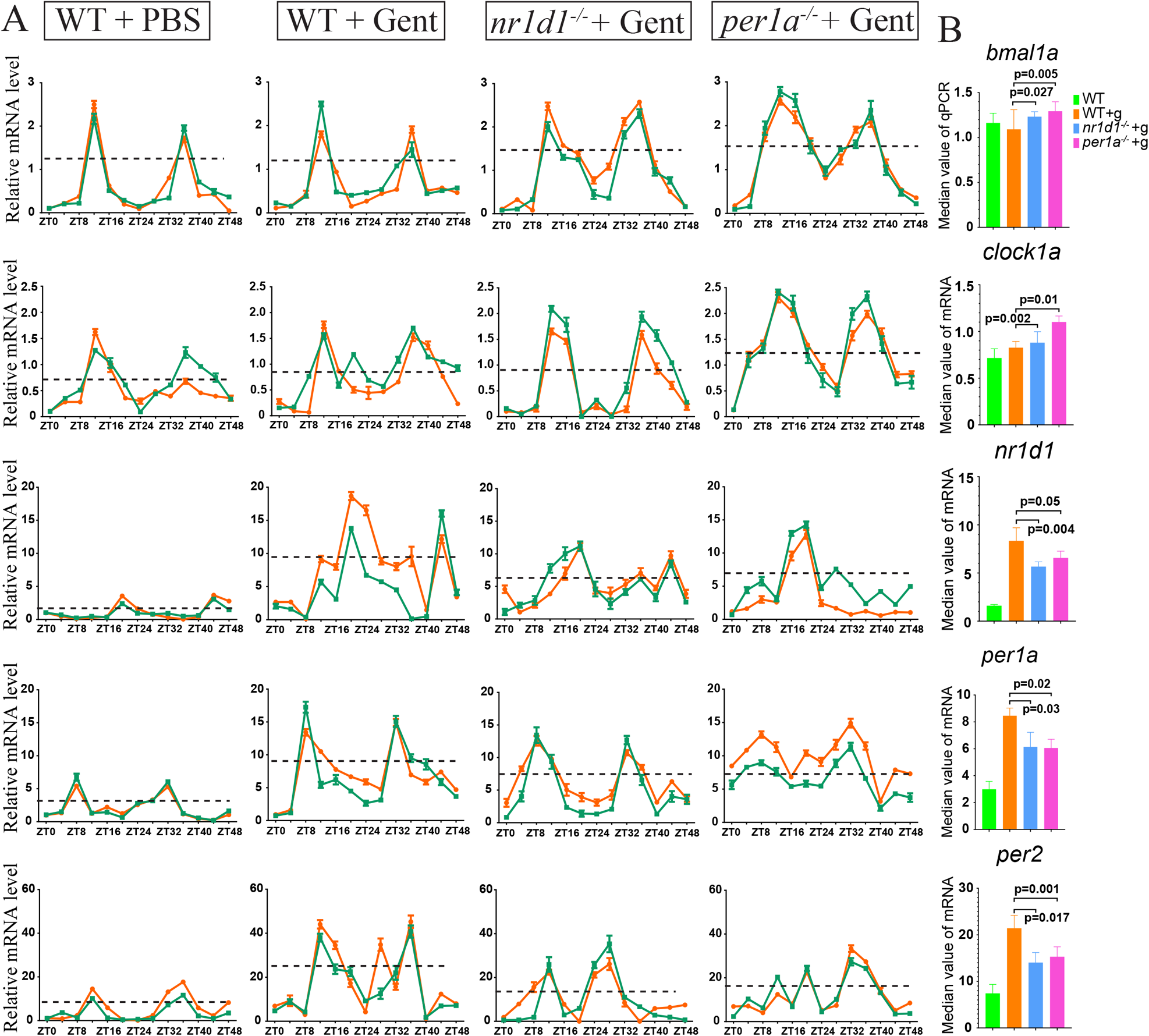
Rhythmic expression of the circadian clock genes. (**a)** qRT-PCR analysis showing the rhythmic expression of the indicated clock genes at 4h time points over 48 hours, with and without gentamicin (Gent) treatment. qRT-PCR were repeated at least three times. The green and red colors were used to represent averaged values of two replicates (each with 3 biological repeats). The dashed black lines indicated the median values of mRNA levels. (**b)** Quantification of panel a, based the median values of mRNA levels. P-values for each panel were indicated to show the statistical significance.

To investigate the function of the clock genes, we generated *per1* and *nr1d1* mutants by using the CRISPR/Cas9 technology.^46^ Two mutant alleles for each gene were identified, as revealed by genotyping (Supplementary Figure 4A). We monitored the temporal expression of the core clock genes in *per1a* and *nr1d1* mutants after gentamicin treatment. Compared to wild type controls, *per1a* and *nr1d1* were significantly down-regulated (*P*< 0.05) (Figure 4A-B).

To explore whether *per1* and *nr1d1* are important for kidney regeneration, renal functional recovery analysis was performed. Approximately 65% and 80% of nephrons in wild type animals were functional at 7 and 10 dpi, as revealed by dextran fluorescent signals. By contrast, the percentage was merely 20% and 40% in mutant kidneys, respectively (Figures 5A-B). HE staining and WISH experiments confirmed that kidney repair was affected or delayed in the absence of *per1* and *nr1d1* (Figures 5C-D; Supplementary Figure 5A).

**Figure 5.**
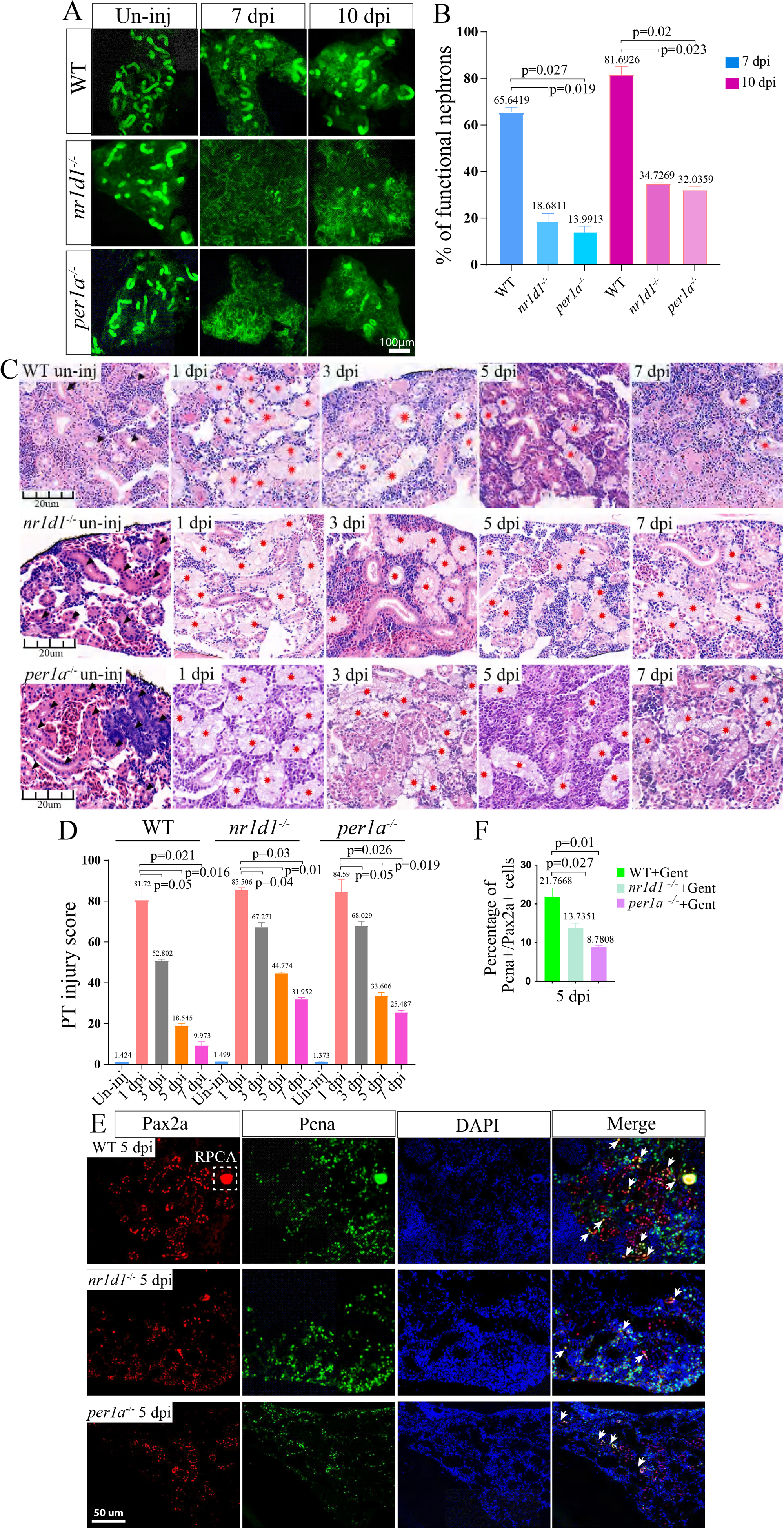
Loss of *per1* and *nr1d1* leads to renal regeneration defects. (**a)** Representative fluorescence signals of kidneys at the indicated time points, in control, *per1a−/−*, and *nr1d1−/−* animals. (**b)** Quantification of panel a (n=3-5 different regions of each group), showing the percentage of functional nephrons. (**c)** HE staining results of kidneys at the indicated time points, in control, *per1a−/−*, and *nr1d1−/−* animals. (**d)** Quantification of panel c (n=4-5 different regions of each group), to show the renal recovery dynamics after AKI. The PT injury score is used to show the percentage of abnormal PTs. (**e)** Pax2a and Pcna double staining images of kidneys from control, *per1a−/−*, and *nr1d1−/−* animals at 5 dpi. White arrows: Pax2a and Pcna double positive single cells. White box with dashed line: Pax2a expressing RPC aggregates. (**f)** Quantification of panel e (n=4-5 different regions of each group), with p-values. Gent: gentamicin. The IF and HE were repeated three times, and shown were representative data. P-values for each panel were indicated to show the statistical significance.

Next, double immunofluorescence (IF) staining was performed using Pax2a and Pcna antibodies. Pax2a is a marker of aggregating RPCs.^1^ The results showed that the average percentage of Pax2a^+^ Pcna^+^ cells in mutant kidneys was merely 1/2 to that in controls (Figure 5E-F). Note that the Pax2a^+^ aggregating RPCs was lower in *per1−/−* and *nr1d1−/−* mutant kidneys than in controls.

The observed phenotypes could be due to exaggerated kidney damage and/or increase in apoptosis by loss of *per1* and *nr1d1*. IF staining was performed using antibodies against Kim1, a renal injury marker (Supplementary Figure 5B). At 1/3 dpi, Kim1 expression was similar between control and mutant kidneys. TUNEL experiments showed that apoptosis was comparable between mutant and control kidneys (Supplementary Figure 5C). Thus, kidney injury and apoptosis were not affected by loss of *per1* and *nr1d1*.

Taken together, we concluded that the renal regeneration is *per1a* and *nr1d1* dependent. Loss of *per1* and *nr1d1* causes a decrease in the number of nephrogenic RPCs, resulting in a delayed kidney repair.

### Identifying *lima1a* as a clock target gene

To investigate the mechanism by which *per1* and *nr1d1* control renal regeneration, bulk RNA-sequencing was performed for 1, 3, 5 dpi kidneys from wild type, *per1a−/−*, and *nr1d1−/−* zebrafish (Supplementary file 5). The differentially expressed genes (DEGs) at each time points were identified. Approximately 2600 DEGs (fold change> 2; *P*< 0.05) were identified at 1 dpi (Supplementary Figure 6A). KEGG of the down-regulated DEGs at dpi 1 showed the enrichment terms such as circadian rhythms and metabolism (Figure 6A; Supplementary Figure 6B).

**Figure 6.**
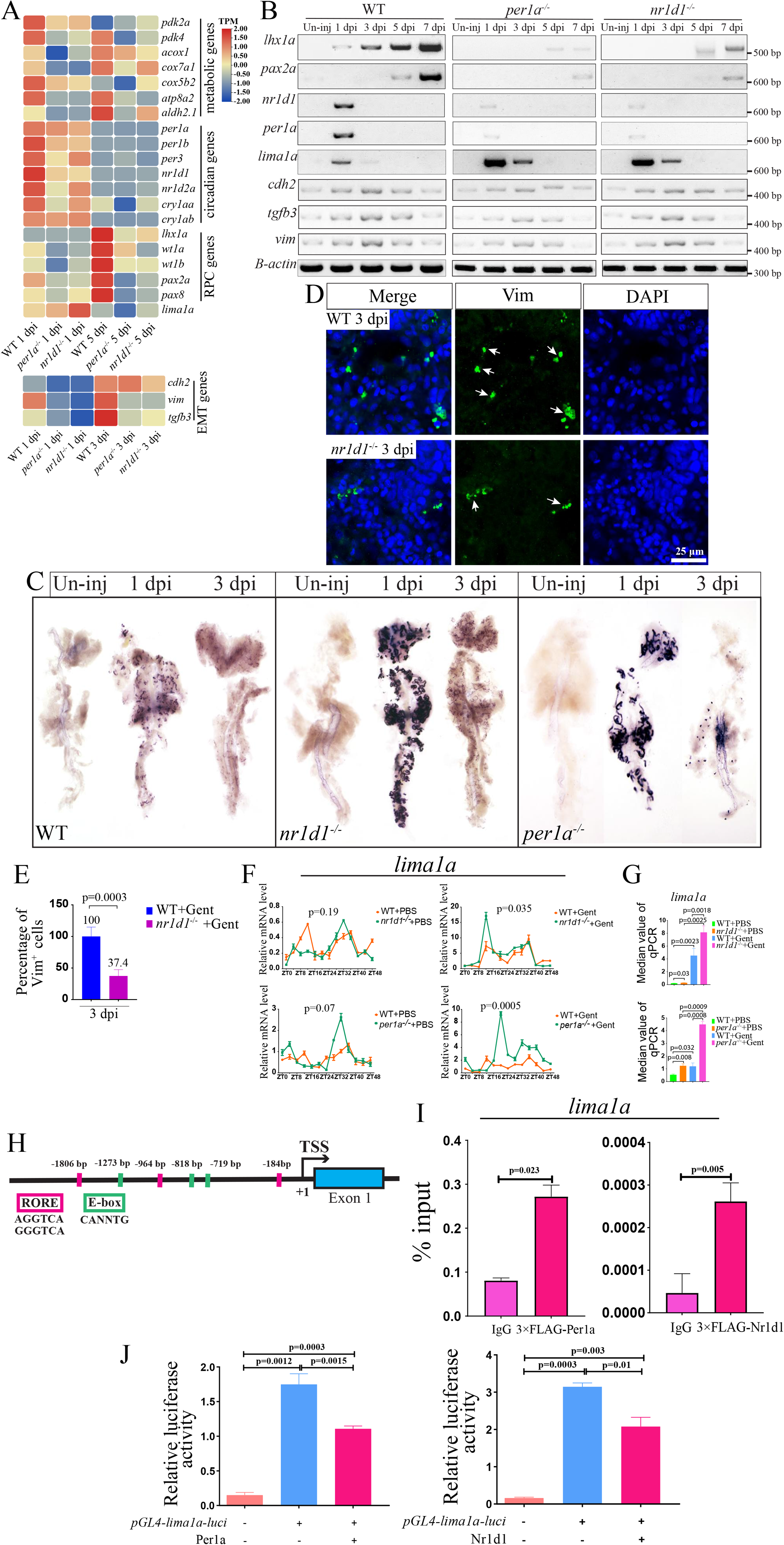
*lima1a* is a downstream target of the clock factors. **(a**) Heat map showing the expression of indicated genes, based on 3 replicates. The mesenchymal genes were placed in the bottom, and other genes were on the top. (b) Gel images of RT-PCR analysis for the indicated genes, in kidneys at the indicated time points of control, *per1a−/−*, and *nr1d1−/−* animals. The RT-PCR were repeated three times, and shown were representative data. (c) WISH results showing the expression of *lima1a* at the indicated time points of control, *per1a−/−*, and *nr1d1−/−* animals. WISH were repeated three times. **(d)** IF staining of Vimentin (Vim) in 3 dpi control and *nr1d1* mutant kidneys. IF experiments were repeated at least two times. **(e)** Quantification of panel d. (**f)** qRT-PCR results showing the rhythmic expression of the indicated clock genes at 4h time points over 48 hours, with and without gentamicin treatment. qRT-PCR were repeated three times, and shown were representative data. (**g)** quantification of panel e. (**h)** Cartoon showing the features of *lima1a* gene promoter. (**i)** ChIP-PCR results showing the enrichment of the indicated proteins at *lima1a* promoter. The ChIP experiments were repeated two times. (**j)** Luciferase assay showing *lima1a* promoter activities. Luciferase experiments were repeated two times. P-values for each panel were indicated to show the statistical significance. ANOVA was used to test the statistical significance, when multiple comparisons were conducted.

Approximately 2800 DEGs (fold change> 2; *P*< 0.05) were identified at 3 or 5 dpi (Supplementary Figure 6A). KEGG of the down-regulated DEGs at dpi 3/5 showed the enrichment terms such as ECM-receptor interaction, and transendothelial migration. Consistently, EMT markers such as *cdh2/N-cad* and *tgfb* were down-regulated in mutant kidneys. Remarkably, RPC markers such as *lhx1a* and *wt1b* were also down-regulated, which was highlighted by the heap map analysis (Figure 6B, Supplementary Figure 6C).

To confirm the RNA-seq results, RT-PCR, IF and WISH experiments were performed. The results confirmed that the expression of mesenchymal and RPC marker genes was decreased in clock gene mutant kidneys compared to controls (Figures 6C-E). We concluded that *per1* and *nr1d1* are required for proper nephron regeneration via regulation of EMT and RPC genes.

The simplest explanation would be that the circadian clock proteins directly control the expression of mesenchymal and RPC marker genes. However, Per1 and Nr1d1 are known to function as a transcriptional repressors in the circadian system, which excludes that the mesenchymal and RPC markers are their direct targets. In searching for candidate genes, *lima1/eplin,* a negative regulators of EMT that is shown to be expressed in kidney tissues, was of great interest to us for a few reasons. First, the expression of *lima1a* was reversely correlated to that of *per1* and *nr1d1* (Figure 6B), and was increased in *per1* and *nr1d1* mutant kidneys (Figure 6C). Second, *lima1a* expression preceded and negatively correlated to that of mesenchymal and RPC marker genes (Figures 6B-D). Third, and importantly, the circadian regulation of cytoskeleton is known to affect wound-healing efficacy.^47^ We speculated that *per1* and *nr1d1* may regulate the formation RPCs or RPC gene expression through regulation of cytoskeleton remodeling or EMT via *lima1a*. To explore this, we first asked whether *lima1a* is rhythmically regulated by Per1 and Nr1d1, by examining its temporal expression patterns. In wild type zebrafish, *lima1a* was moderately expressed, displaying a rhythmic expression pattern, with a peak at ZT12. After gentamicin treatment, *lima1a* was induced in both wild type and clock mutant kidneys, and the phase of circadian rhythm was comparable. However, *lima1a* levels were much higher in clock mutant kidneys than in controls (Figures 6F-G).

Examination of DNA sequences upstream the TSS of the *lima1a* gene revealed the presence of multiple E-box and RORE sites (Figure 6H; Supplementary Figures 6D-E), suggesting that Per1a and Nr1d1 suppress *lima1a* by directly binding to its promoter. To investigate this, we injected mRNAs encoding 3×FLAG-tagged Nr1d1 or Per1 into 1-cell stage of zebrafish embryos, and performed chromatin immunoprecipitation (ChIP) experiments with FLAG antibodies. The results showed that FLAG-Nr1d1 and FLAG-Per1 can occupy *lima1a* promoter (Figure 6I; Supplementary Figure 6D-G).

Next, we made the pGL4-*lima1a*Luc reporter construct, and transfected it with plasmids encoding Nr1d1 and Per1 into HEK293T cells. The luciferase activities were measured, and the results showed that overexpression of Nr1d1 and Per1 decreased *lima1a* promoter activities (Figure 6J, Supplementary Figure 6E). Taken together, we propose that *lima1a* is a direct target of Per1 and Nr1d1.

### Lima1a inhibits renal regeneration

To study the functional role of *lima1a*, we generated zebrafish mutant lines using the CRISPR/Cas9 technology. We designed gRNAs against the exon 2 of the *lima1a* gene. Two mutant alleles were identified, which were confirmed by genotyping (Figure 7A). The mutations led to a truncated protein of 98 aa without functional domains, resulting in loss of function of the Lima1a protein.

**Figure 7.**
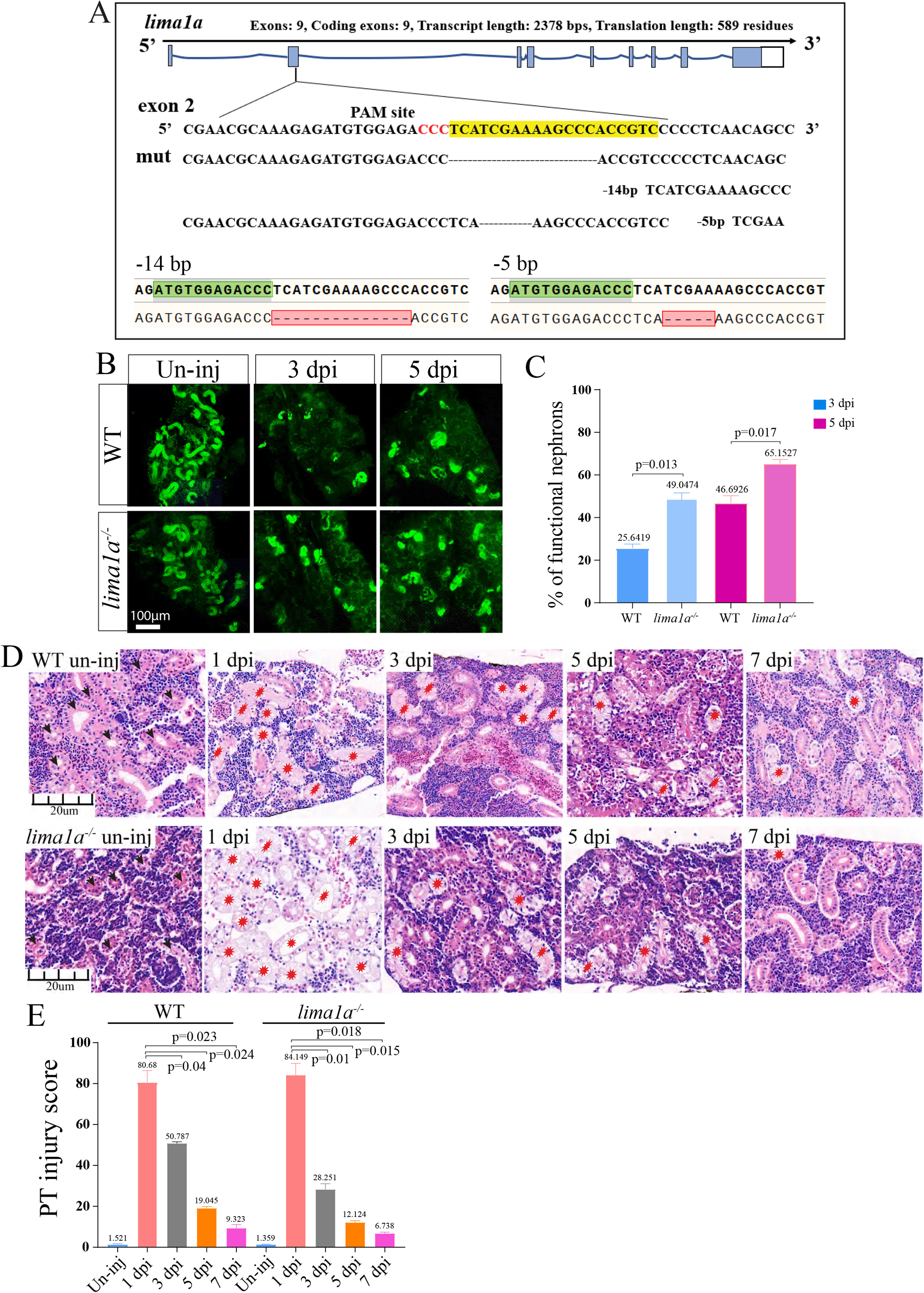
*lima1a* loss accelerates nephron regeneration. (**a)** CRISPR/cas9 design and mutant alleles for *lima1a*. (**b)** Representative fluorescence signals of kidneys at the indicated time points, in control and *lima1a−/−* animals. The experiments were repeated three times, and shown were representative images. (**c)** Quantification of panel b (n = 3-5 different regions of interest per group), showing the percentage of functional nephrons. (**d)** HE staining images of kidneys at the indicated time points, in control and *lima1a−/−* animals. (**e)** Quantification of panel d (n = 4-5 different regions of interest per group), to show the renal recovery dynamics after AKI. The PT injury score is used to show the percentage of abnormal PTs. HE staining were repeated three times, and shown were representative images. P-values for each panel were indicated to show the statistical significance.

The maternal zygotic *lima1a* mutant embryos were morphologically normal, and can be raised up to adults with normal kidney morphology. Renal functional recovery assay was performed for 5 month control and *lima1a−/−* zebrafish following gentamicin treatment. We found that nephron regeneration was enhanced in the absence of Lima1a. At 3 dpi, there were few fluorescent nephrons in control kidneys, by contrast, approximately 20% of nephrons were filled with fluorescent dextran in mutant animals (Figures 7B-C; Supplementary Figure 7A-B). Similar results were observed at 5 dpi. HE staining confirmed that nephrons were better repaired in 3/5 dpi mutants than in controls (Figures 7D-E). Based on the data, we concluded that *lima1a* is a negative regulator of renal regeneration, and its loss accelerates regeneration process.

### *lima1a* controls RPC genes via EMT

To explore how *lima1a* is involved in renal regeneration, bulk RNA-sequencing was performed for 0, 1, 3, 5 dpi kidneys from wild type and *lima1a* mutant zebrafish (Supplementary file 6), and the DEGs at each time points were identified. Approximately 2000 DEGs (fold change> 2; *P*< 0.05) were identified at 1 dpi (Supplementary Figure 8A). GO enrichment of the up-regulated DEGs at 1 dpi showed the terms such as circadian rhythm, and stress response (Figure 8A). Consistently, circadian genes *per1* and *nr1d1*, early response genes *atf3* and *egr* were up-regulated (Figure 8B). Approximately 1800 and 2900 DEGs (fold change> 2; *P*< 0.05) were identified at 3 and 5 dpi, respectively (Supplementary Figure 8B). GO of the up-regulated DEGs at 3 and 5 dpi showed the enriched terms such as actin nucleation, regulation of actin cytoskeleton, and extracellular matrix organization. Specifically, mesenchymal and cytoskeletal genes such as *cdh2* and *vim* were up-regulated.

**Figure 8.**
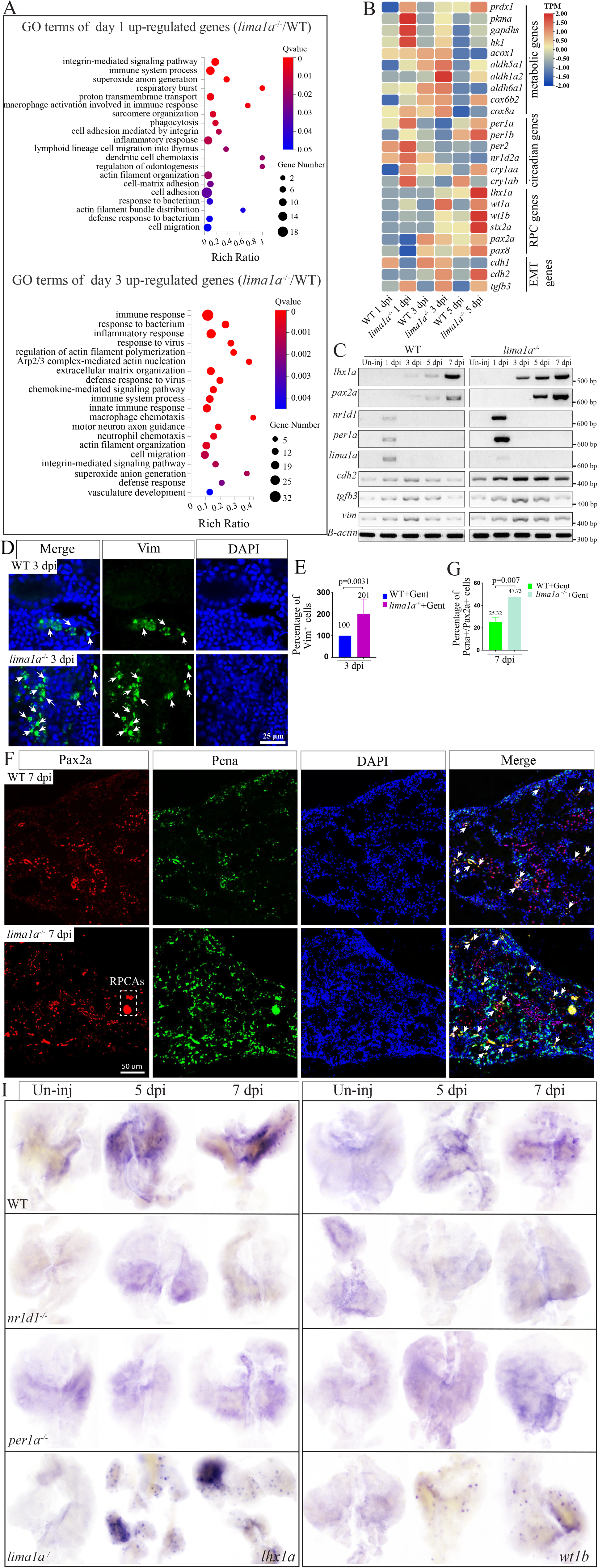
*lima1a* controls RPC genes via EMT. (**a)** GO terms of up-regulated genes between control and *lima1a−/−* kidneys at 1 dpi. (**b)** Heat map showing the expression levelsof the indicated genes at the indicated time points, based on three replicates. (**c)** Gel images of RT-PCR products showing the dynamic expression of the indicated genes at the indicated time points. The experiments were repeated three times and shown were representative. (**d)** IF staining images showing the expression of Vimentin in control and *lima1a−/−* kidneys at the indicated time points. The experiments were repeated two times and shown were representative. **(e)** Quantification of panel d. (**f)** IF staining images showing the expression of Pax2a and Pcna in control and *lima1a−/−* kidneys at 7 dpi. White dashed box: Pax2 expressing RPC aggregates. The experiments were repeated three times and shown were representative. (**g)** Quantification of e (n =4-5 different regions of interest per group). (**h)** WISH images showing the expression of the indicated RPC markers at the indicated time points in control and mutant kidneys. The experiments were repeated three times and shown were representative. P-values for each panel were indicated to show the statistical significance.

As hallmarks of EMT include concomitant downregulation of epithelial E-Cadherin (*cdh1*) and upregulation of N-cadherin (*cdh2*) and *vimentin*, our RNA-seq data suggested that *lima1a* is a regulator of EMT in zebrafish kidneys.^18, 48^ To further investigate this, RT-PCR and IF experiments were performed (Figure 8C-F). The results confirmed that *lima1a* is a negative regulator of EMT, and its loss leads to increased expression of mesenchymal markers.

Importantly, the expression of RPC markers was markedly increased, along with that of mesenchymal markers in *lima1a* mutant kidneys (Figure 8C-H; Supplementary Figures 8C-D), explaining why loss of *lima1a* accelerates the nephron repair process. Interestingly, the expression of *nr1d1* and *per1* became much higher in *lima1a* mutant kidneys than in controls, indicating that *lima1a* and *nr1d1/per1* antagonize each other. Taken together, we concluded that *lima1a* is involved in the regulation of RPC markers or the formation of RPCs via modulation of EMT, an essential biological process required for kidney regeneration.^15^

## DISCUSSION

A recent study in mice has shown that the core clock genes such as *Per2* are required for embryonic kidney development. However, its role in kidney regeneration has not been well defined. By analysis of our scRNA data, we found that clock genes such as *per1* and *nr1d1* were highly induced in PTECs after AKI. The observation suggested that the circadian clock genes might be involved in renal regeneration. By generating clock gene mutant zebrafish, we provided compelling evidence that the clock genes are important for proper renal regeneration by control of EMT of the surviving PTECs via suppressing *lima1a*, thereby facilitating the expression of RPC markers such as *lhx1a* and *wt1*.

It is now widely accepted that renal regeneration occurs predominantly from tubular epithelial cells that reside within the injured kidney, with minimal contribution from extra-renal cells.^1,45^ Therefore, although we identified a *sall1* and *cdkn*-expressing renal stem/progenitor cell sub-population by scRNA analysis, we primarily focused on PTECs. We surprisingly found that RPC markers such as *lhx1a* and *wt1b* were highly induced within PTECs at 5 dpi. This is accompanied (preceded) by increased expression of mesenchymal markers (concomitant with decreased expression of epithelial markers). Based on these observations, we hypothesized that the mesenchymal-like *lhx1a*^+^ RPCs might be derived from tubular epithelial cells through EMT. In fact, previous work has suggested that EMT is involved in the formation and migration of *lhx1a^+^* RPCs.^3^ We propose that Lima1, a cytoskeleton binding protein that functions to negatively regulate epithelial to mesenchymal transition, plays a key role in this process. A previous work has shown that Lima1 protein is expressed in renal PTECs and glomeruli cells, and modulate cell adhesion and movement.^19^ By generating *lima1a* mutants, we demonstrated that renal regeneration process was accelerated, concomitant with increased expression of EMT- and RPC markers. Our results thus support a scenario, where the surviving PTECs acquire stem cell-like properties during dedifferentiation and EMT to become *lhx1a^+^*RPCs, which then migrate to regenerate the nephrons.

We found that *per1* and *nr1d1* are induced in PTECs, and required for proper renal regeneration. In *per1* and *nr1d1* mutant kidneys, mesenchymal and RPC marker genes were down-regulated, which strongly indicated that the clock genes contribute to nephron regeneration by regulation of EMT and the formation of RPCs. Per1 and Nr1d1 are transcriptional repressors of the circadian system, excluding that they directly regulate the mesenchymal and RPC marker genes. By RNA-seq, ChIP, and IF analyses, we identified *lima1a* as the direct target of the circadian clock proteins. The circadian regulation of cytoskeleton is known to affect wound-healing efficacy in skin.^47^ Our work may provide general mechanistic explanation of how the circadian clock regulates actin cytoskeleton remodeling for tissue regeneration.

In summary, we show that circadian regulation of the cytoskeleton of tubular epithelial cells is important for renal regeneration, through control of the generation and proliferation of *lhx1a^+^* RPCs. Our work has three key findings. First, the *lhx1a^+^* RPCs can be derived from surviving PTECs via EMT. Second, circadian regulation of actin cytoskeleton is important for kidney regeneration via *lima1a,* providing evidence how circadian regulation of cytoskeleton is achieved. ^47^ Third, circadian clock genes promote renal regeneration by regulating the formation of RPCs. Our data revealed how the circadian clock genes control the expression of RPC genes by regulation of dedifferentiation and EMT of intrinsic TECs through remodeling of actin cytoskeleton.

This work was primarily performed using the zebrafish model. And, exactly how night/day cycles impact kidney repair remains to be investigated. Moreover, as the work was designed to focus on the hypothesis that injured TECs acquire stem cell-like properties and contribute to nephron repair, the type of nephron regeneration (stem cell neonephrogenesis vs repair of existing tubules) was not discussed. In the future, it will be important to investigate these in more detail, by using mammalian models such as mouse or rat.

## Data sharing statement

The bulk and scRNA-seq data have been deposited in China National Center for Bioinformation (CNCB) in Beijing at https://www.cncb.ac.cn/. The accession number of scRNA-seq data is CRA014875: https://bigd.big.ac.cn/gsa/browse/CRA014875. The accession numbers of bulk RNA-seq data are CRA014877: https://bigd.big.ac.cn/gsa/browse/CRA014877, and https://bigd.big.ac.cn/gsa/browse/CRA014881. Other data are available upon reasonable request and free for sharing.

## Acknowledgement

This work was supported by National Key Research and Development Program of China (2022YFA0806600) and the Strategic Priority Research Program of the Chinese Academy of Sciences (XDB31000000). We thank Dr. Yong Long from IHB for providing *nr1d1* mutant zebrafish.

## Supplementary information

The supplementary methods, tables and figures can be found in the supplementary files.

## Supplementary Figures and legend

**Supplementary Figure 1.**
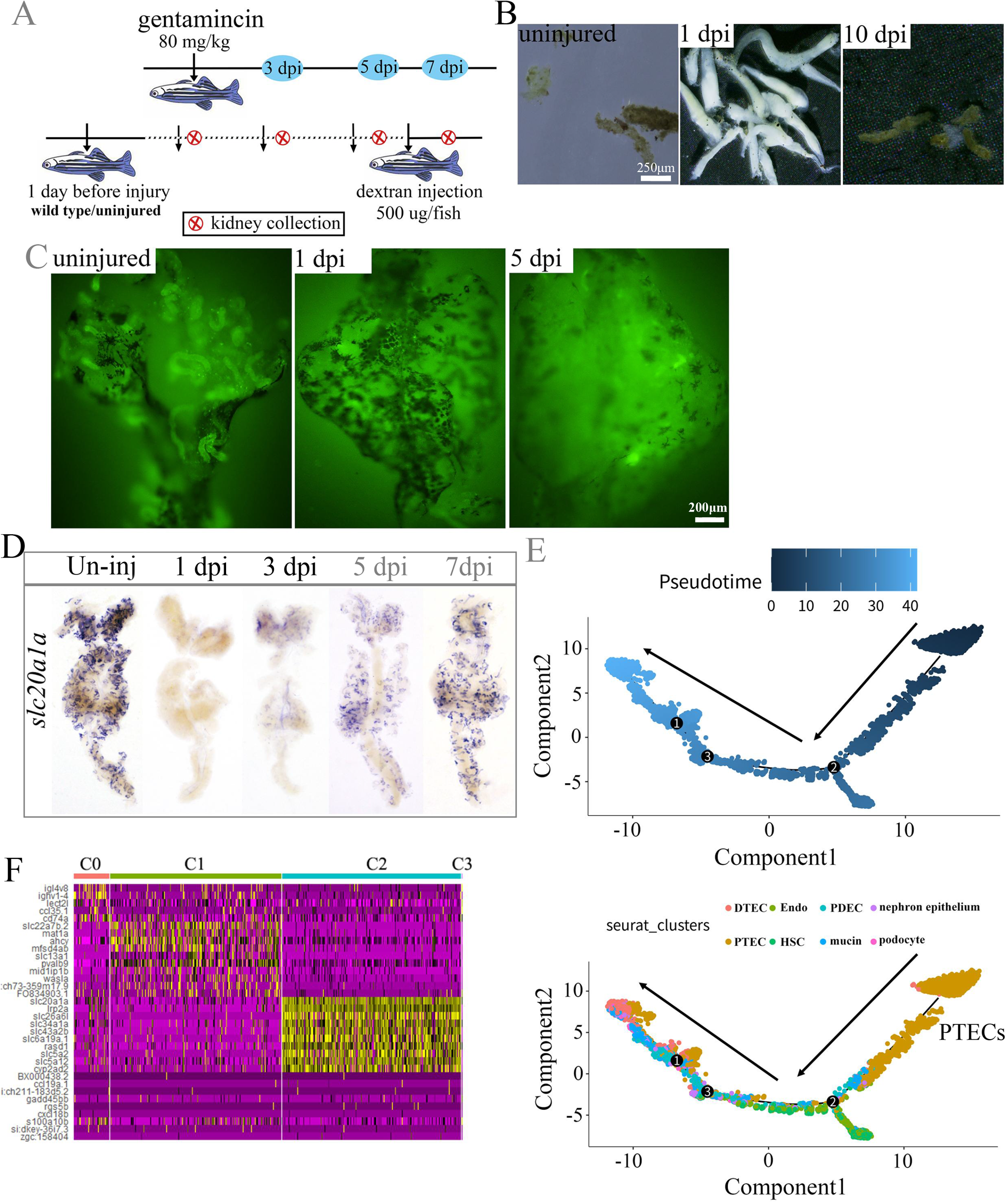
Gentamicin treatment leads to acute kidney injury. **(a)** Diagram showing additional information of experimental design. (**b)** Images showing the casts of dead epithelial tissues in the indicated time points. **(c)** Fluorescence signal at the indicated time points. **(d)** WISH images showing the dynamic expression of *slc20a1a* at the indicated time points. WISH experiments were repeated 3 times, and shown were representative. (**e**) Pseudotime analysis showing that PT cluster is at the start of the trajectory. **(f)** Heat map showing the top enriched genes in the 4 subclusters.

**Supplementary Figure 2.**
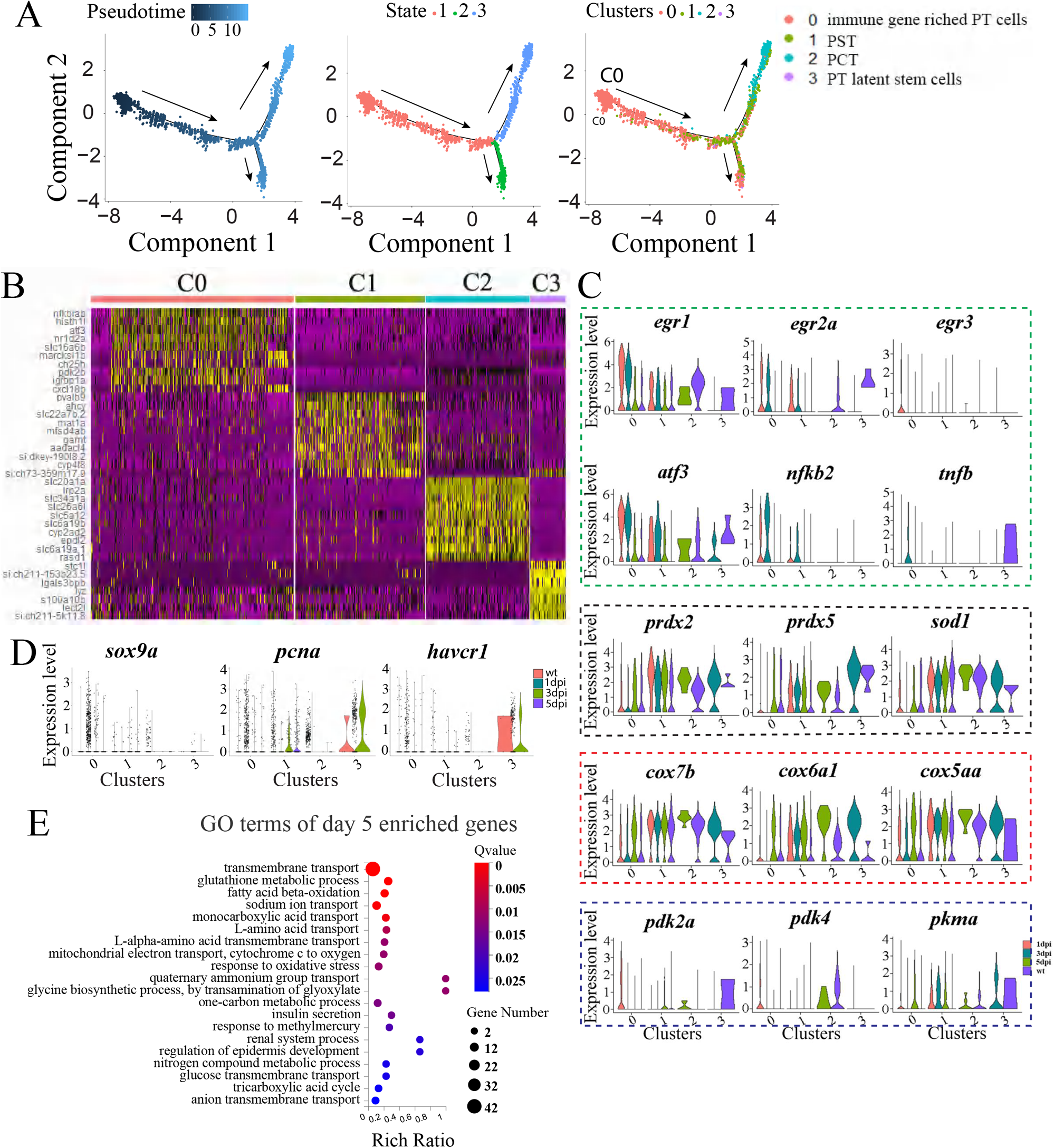
Dynamic gene expression of the 4 PT subclusters. **(a)** The top enriched genes of the 4 subclusters after AKI. **(b)** Violin plots showing the dynamic expression of the indicated genes, based on analysis of scRNA-seq data. The dashed colored boxes were used to group the indicated markers, including early response-, anti-oxidant-, and metabolic-genes. **(c-d)** Violin plots showing the dynamic expression of the indicated genes, based on analysis of scRNA-seq data. **(e)** GO terms of day 5 DEGs.

**Supplementary Figure 3.**
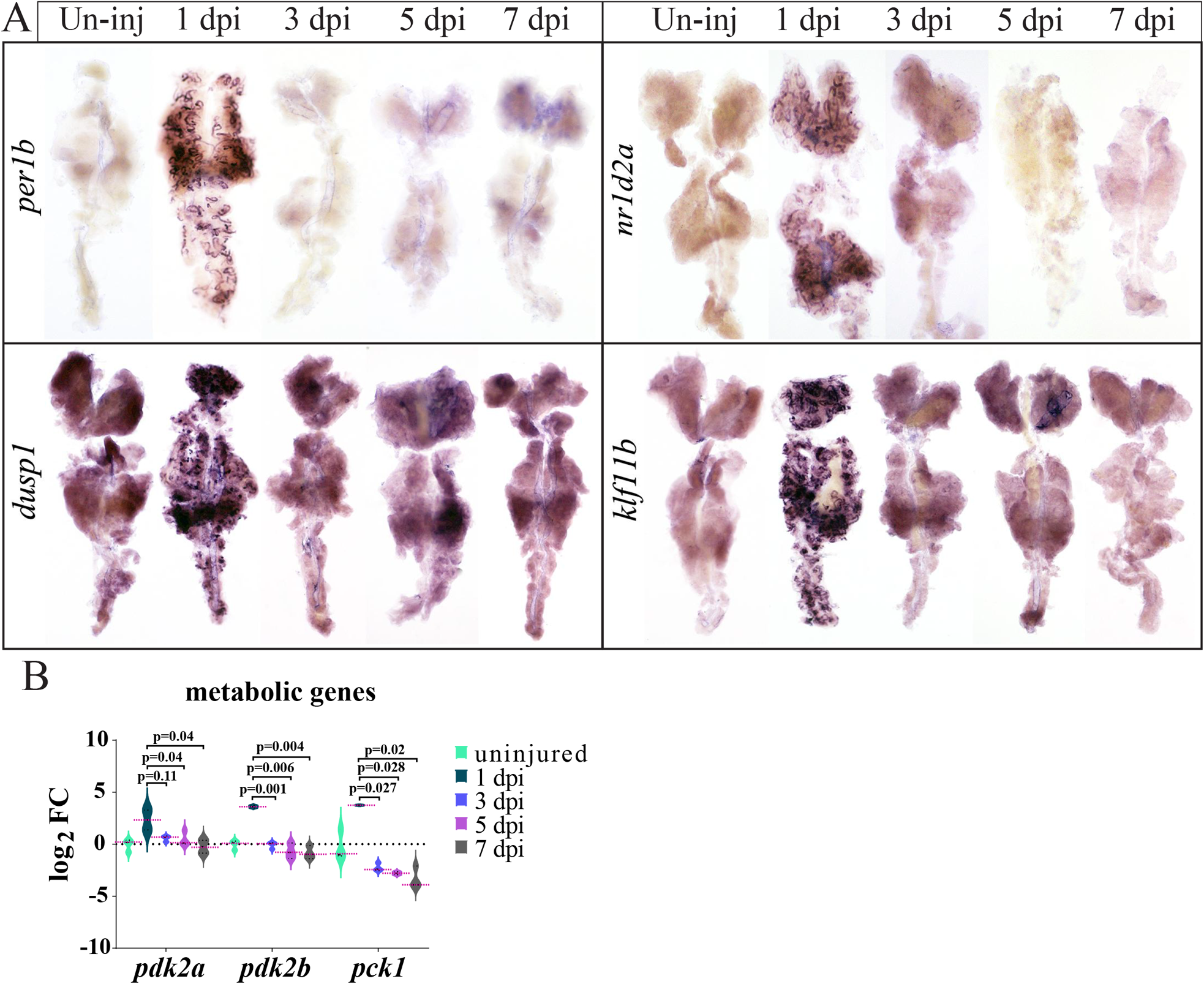
Dynamic expression of indicated genes in response to gentamicin. **(a)** WISH images showing the dynamic expression of the indicated marker genes. **(b)** Dynamic expression of the indicated genes, based on bulk RNA-seq data. P-values were indicated to show the statistical significance.

**Supplementary Figure 4.**
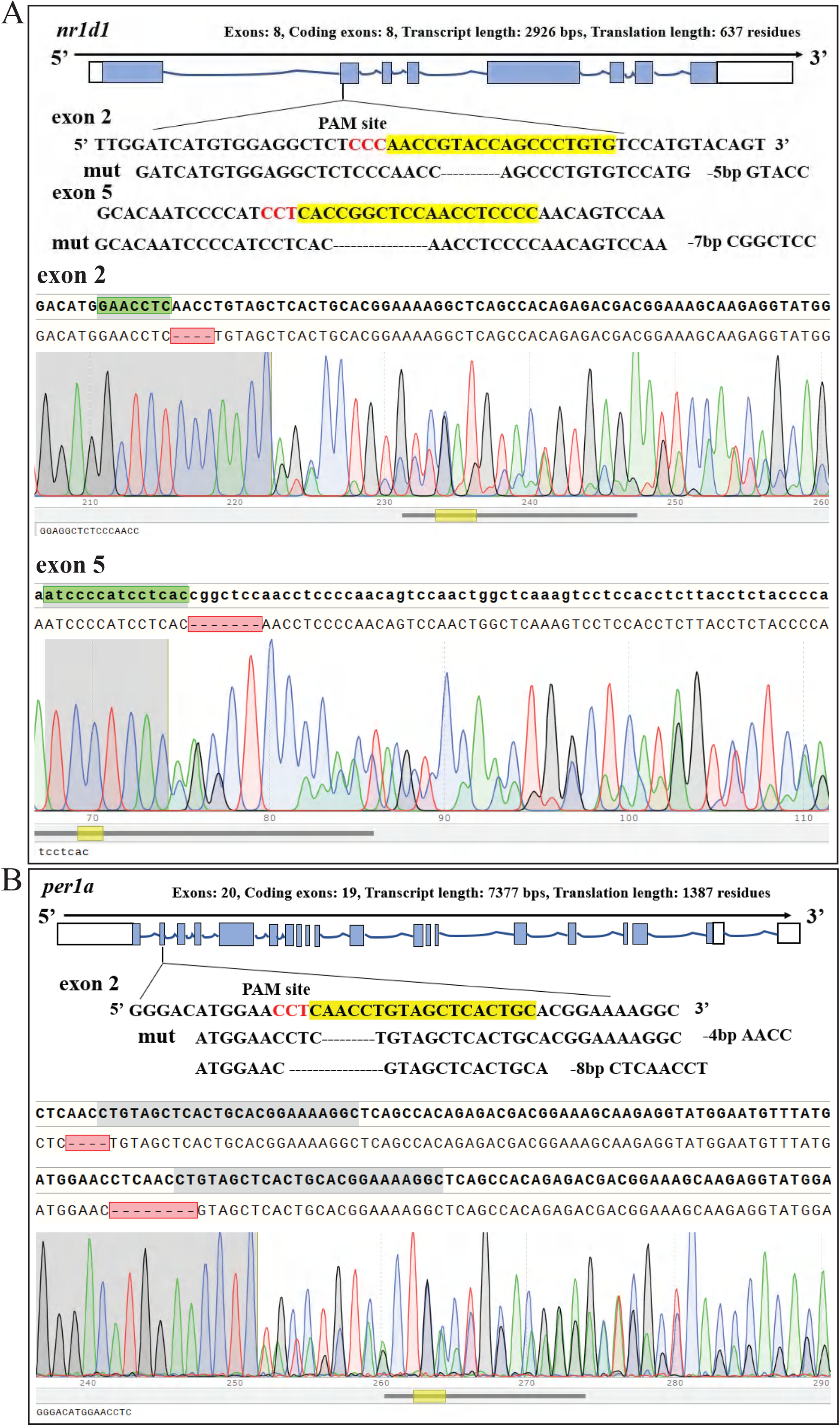
Generation of clock gene mutants by CRISPR/cas9. **(a)** Design of gRNAs for *nr1d1* and the genotyping results. **(b)** Design of gRNAs for *per1a* and the genotyping results.

**Supplementary Figure 5.**
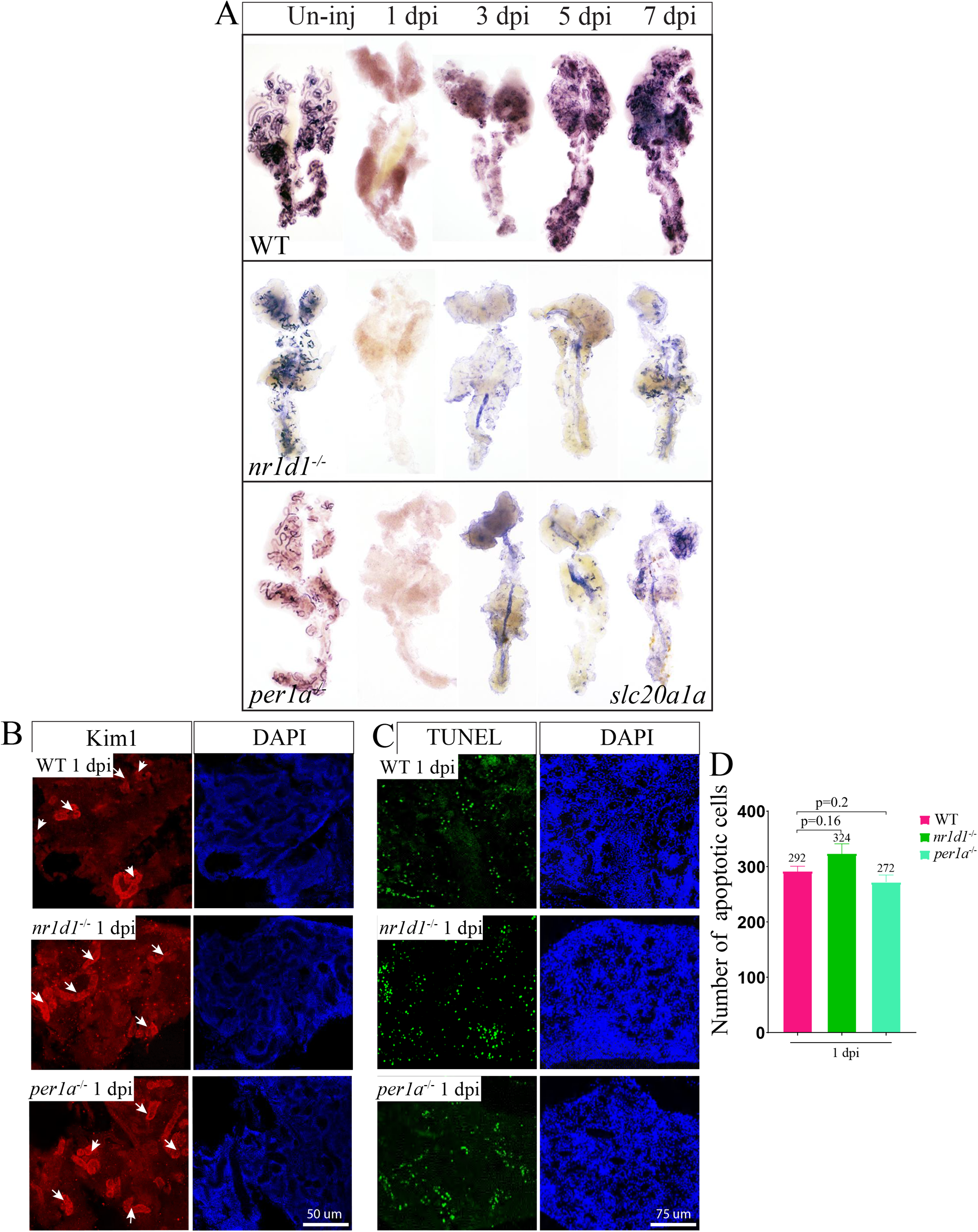
Expression of slc20a1a. **(a)** WISH images showing the expression of *slc20a1a* at the indicated time points, in control and mutant kidneys. **(b)** IF staining of Kim1 of kidneys from control, *per1a−/−*, and *nr1d1−/−* animals, at the indicated time points. (**c)** TUNEL results of kidneys at the indicated time point, in control, *per1a−/−*, and *nr1d1−/−* animals. (**d)** quantification of panel c.

**Supplementary Figure 6.**
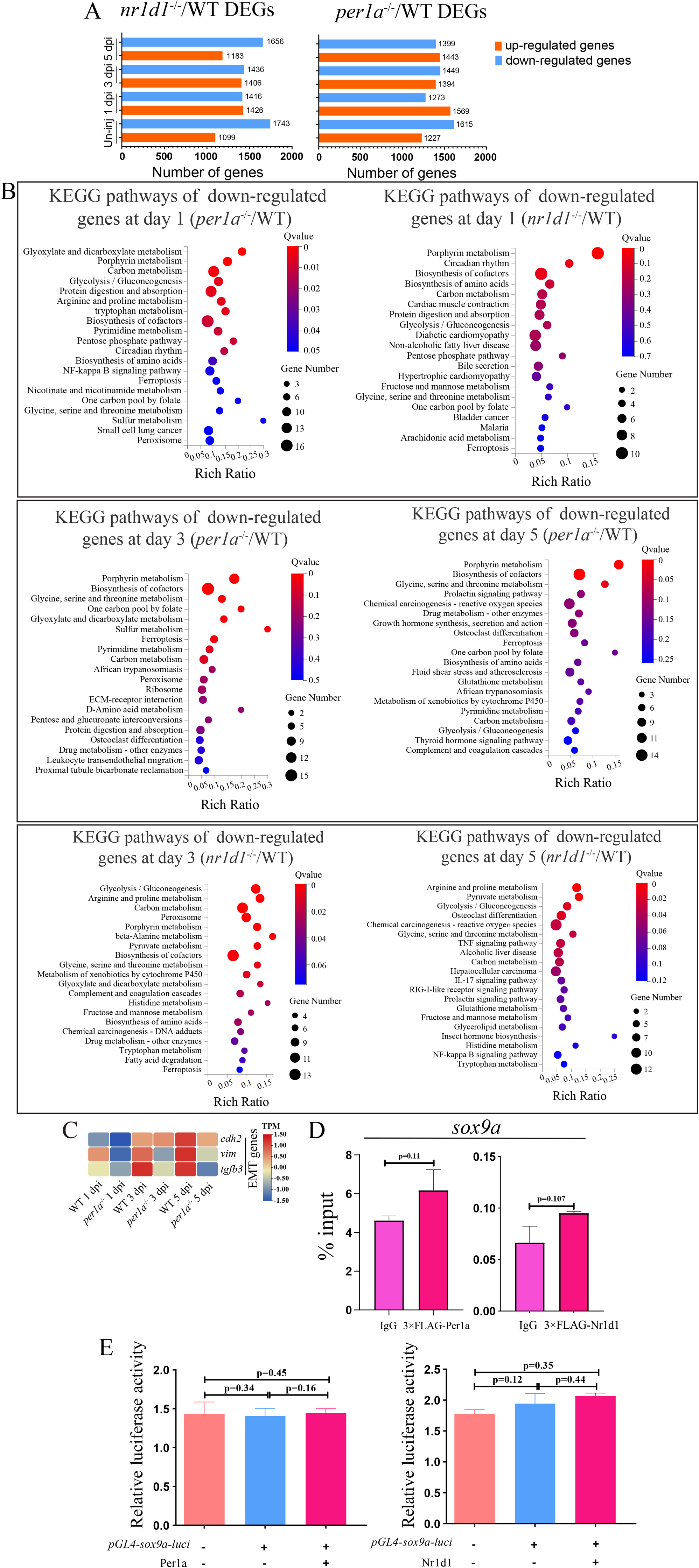
*lima1a* is negatively regulated by the clock factors. **(a)** Up- and down-regulated DEGs between control and clock gene mutant kidneys, at the indicated time points. **(b)** KEGG terms of the DEGs at the indicated time points. **(c)** Heat map showing the dynamic expression of EMT marker genes in control and *per1* mutants. **(d)** ChIP data showing the enrichment of FLAG-tagged Per1 and FLAG-tagged Nr1d1 at *sox9a* promoter. **(e)** Luciferase assay showing the effects of expression of FLAG-tagged Per1 and FLAG-tagged Nr1d1 on *sox9a* promoter activities, in HEK293T cells. P-values for each panel were indicated to show the statistical significance. ANOVA was used to test the statistical significance, when multiple comparisons were conducted.

**Supplementary Figure 7.**
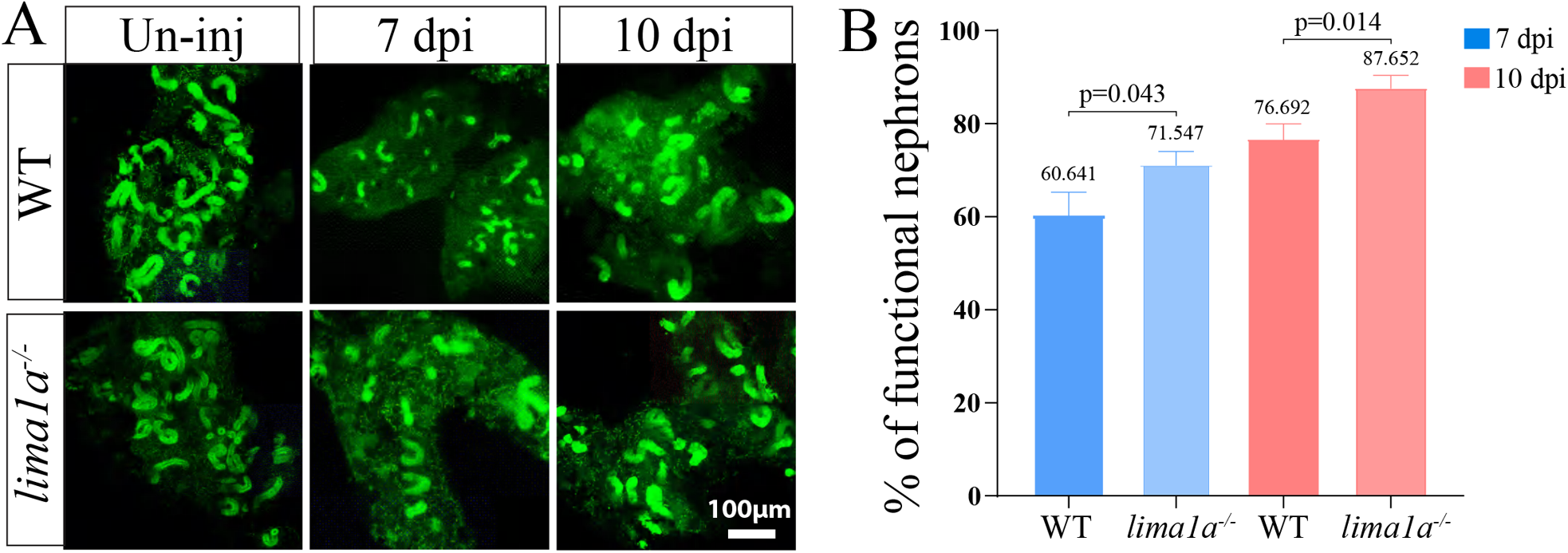
*lima1a* knockout accelerates kidney repair. **(a)** Dextran fluorescent signals in control and *1ima1a* mutant kidneys, at 7 and 10 dpi. **(b)** quantification of panel a (n=4-5 different regions of each group), showing the percentage of functional nephrons.

**Supplementary Figure 8.**
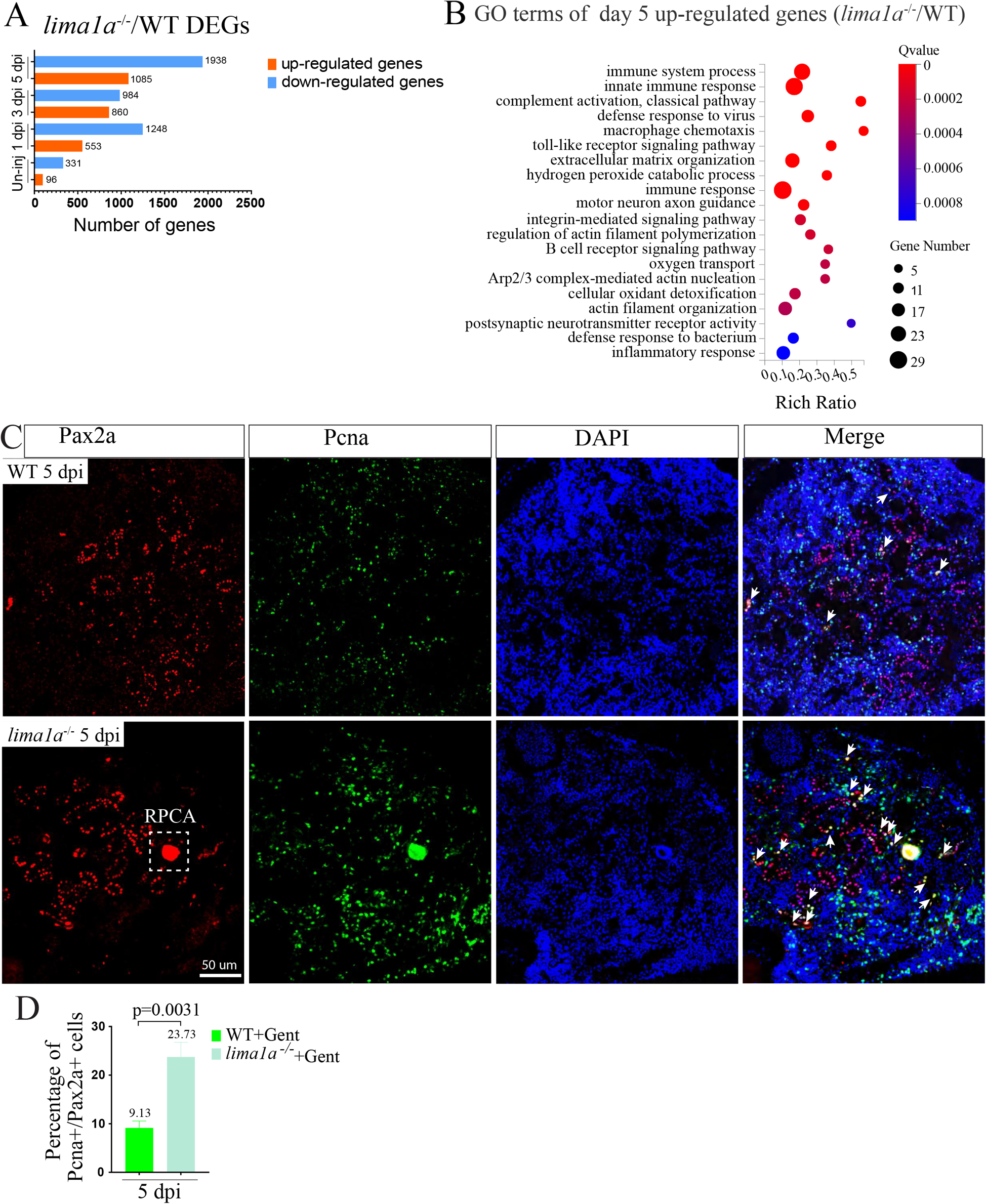
Analysis of DEGs between control and *lima1a* mutant kidneys. **(a)** Up- and down-regulated DEGs between control and *lima1a* gene mutant kidneys, at the indicated time points. **(b)** GO terms of the up-regulated DEGs at the indicated time points. **(c)** IF staining images showing the expression of Pax2a and Pcna in control and *lima1a−/−* kidneys at the indicated time point. White dashed box: Pax2 expressing RPC aggregates. The experiments were repeated three times and shown were representative. **(d)** quantification of c (n=4-5 different regions per group).

